## Supplemental Materials for "A BioID-derived proximity interactome for SARS-CoV-2 proteins"

**Supplementary Materials**

**Table S1. Addgene constructs and primers used for SARS-CoV-2 BioID fusion-proteins.**

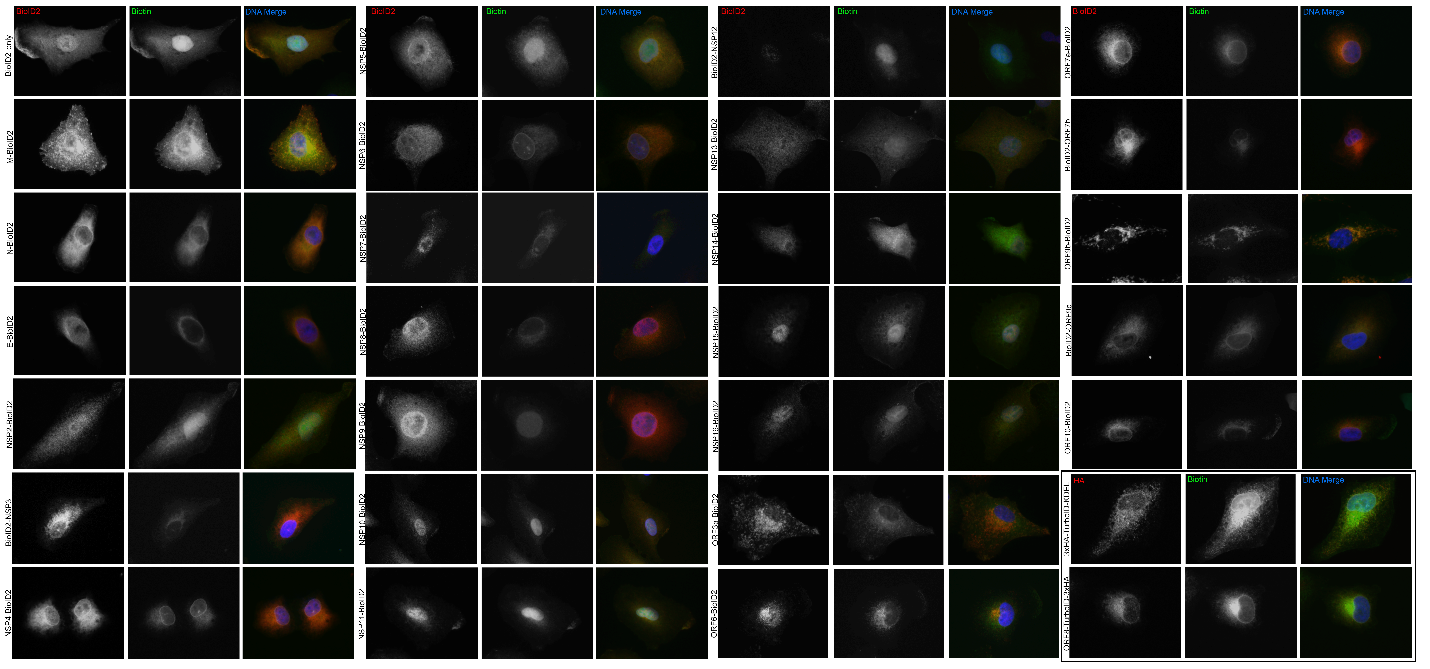

**Figure S1. Epifluorescent images of all BioID fusion-protein localizations in A549 cells.** A549 human lung cells stably expressing SARS-CoV-2 BioID2 fusion-proteins were assessed for fusion-protein expression and localization (red) and promiscuous biotinylation (green) following the addition of exogenous biotin.

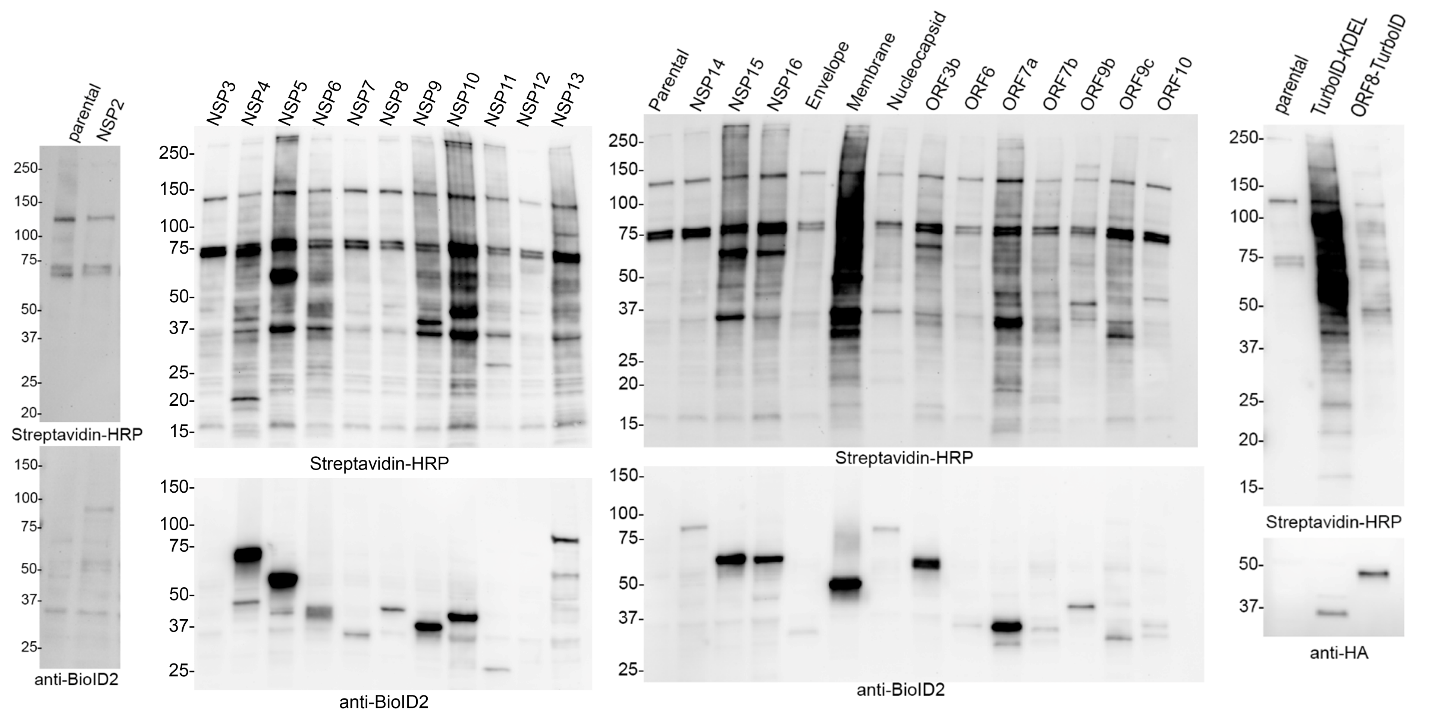

**Figure S2. Western blot analysis of A549 cell stably expressing SARS-CoV-2 BioID fusion proteins.** A549 cells stably expressing BioID2 fusion-proteins were evaluated for fusion-protein expression (anti-BioID2) and promiscuous biotinylation (streptavidin-HRP) and A549 cells stably expressing TurboID-KDEL control or ORF8-TurboID were evaluated for fusion-protein expression (anti-HA) and promiscuous biotinylation (streptavidin-HRP).

**Table S2. Global proteomic analysis of significantly up- and down-regulated proteins in response to viral protein expression.**

**Table S3. SARS-CoV-2 viral-host interaction candidates**

**Table S4. Overlap with the host restrictome.**

**Table S5. High-confidence viral-host interactions**
